# Insecticide resistance in *Aedes aegypti* from the National Capital Region, Philippines

**DOI:** 10.1101/2023.09.08.556786

**Authors:** Jason R. Angeles, Richard Paul B. Malijan, Ariza Minelle A. Apilado, Mary Ann T. Ammugauan, Ferdinand V. Salazar

**Author notes:** These authors contributed equally to this work (Jason R. Angeles, Richard Paul B. Malijan).

## Abstract

**Background:** Human arboviral diseases such as dengue, chikungunya and Zika can be transmitted by the mosquito *Aedes aegypti*. The insecticide-based vector control strategy is critical in reducing the transmission of these *Aedes*-borne diseases but is threatened mainly by the emergence of insecticide resistance.

**Methodology/Principal Findings:** Adult *Aedes aegypti* from National Capital Region, Philippines were subjected to bioassay to determine their susceptibility to the diagnostic doses of pyrethroid, organochlorine and organophosphate insecticides following the standard World Health Organization insecticide susceptibility test. This study reports for the first time the existence of insecticide resistance in *Ae. aegypti* from the Philippines to pyrethroids and organochlorine. Results from this study showed that most of the *Ae. aegypti* populations exhibited phenotypic resistance to the pyrethroids (permethrin and etofenprox) and an organochlorine (DDT) while all populations tested to malathion were still susceptible to this organophosphate. Varying resistance levels to deltamethrin, cyfluthrin and lambdacyhalothrin were also observed in the different mosquito populations.

**Conclusions:** Insecticide resistance exists in local populations of *Ae. aegypti* from the National Capital Region. This finding should alert public health authorities to consider modifying the existing vector management package for greater control efficacy. Best practices that are proven to prevent and/or delay the development of insecticide resistance such as insecticide rotation should be implemented. Alternative toxicants and chemicals with a different mode of action, such as repellents, should be explored to ensure continuing efficacy of program interventions.

**Author summary:** The National Capital Region (NCR), Philippines reports the country’s highest dengue incidence. Apart from being populous and the center of economic activity, the local government authorities of this region have undertaken significant vector control efforts devoted to dengue. The use of insecticides to reduce mosquito vector density remains the handiest control method. This scenario necessitated the documentation of the resistance levels, particularly of the most important vector *Aedes aegypti*. An insect is said to be resistant when the known effective dose of an insecticide can no longer sufficiently kills the same insect population. This study showed that *Ae. aegypti* population from cities in NCR had developed resistance to commonly used pyrethroids (permethrin, etofenprox) and to an organochlorine (DDT). Highly localized variations of resistance and susceptibility within cities at NCR were recorded against deltamethrin, cyfluthrin and lambdacyhalothrin. This finding should alert public health authorities to consider modifying the existing vector management package for greater control efficacy.

## Introduction

The public health implication of dengue infection in the Philippines has been significant since its first recorded outbreak [1]. The high numbers of reported cases in previous years showed that it has been causing damaging effects on human lives. In 2019, the Philippine Department of Health (DOH) declared a National Dengue Epidemic due to the significantly higher cases than in previous years [2]. However, the impact of community lockdowns to contain the COVID-19 pandemic has dropped national dengue figures significantly [3,4]. This situation supports earlier claims that routine house-to-house human movements aid the spread of dengue in localities [5]; much so that human travel on both local and global scales, represents a significant global health risk, particularly in areas with changing climatic suitability for the mosquito vector [6].

Human arboviral diseases such as dengue, chikungunya and Zika remain a global public threat even in the advent of the worldwide coronavirus pandemic [7,8]. These diseases are transmitted by *Aedes aegypti*; a highly adaptive species that thrives both in urban and suburban areas. There is no specific prophylactic treatment nor readily accessible vaccine for *Aedes*-borne diseases to date. The future of vaccines against dengue remains under litigation due to perceived side effects after the initial mass administration resulting in citizens’ hesitancy [9]. Thus, there remains a need to develop and implement vector control measures to combat such diseases.

Insecticide use has been the cornerstone of *Aedes* vector control in the Philippines. The use of safe and efficacious insecticides against the adult and larval populations of mosquito vectors is one of the most effective ways to rapidly interrupt the transmission of mosquito-borne diseases [10]. The high efficacy in regulating the mosquito populations with relatively rapid action of insecticide application makes it the most extensively practiced control of *Ae. aegypti* [7]. Utilization of space-spray techniques such as thermal fogging and ultra-low volume (ULV) spraying for control of *Ae. aegypti* adults result in immediate kill effects that drastically lower mosquito populations after the declaration of an epidemic [11,12]. However, with greater coverage and regular use for mosquito control, there is a higher potential risk for vector mosquitoes to develop some form of resistance. The steep increase in insecticide-based intervention means increased exposure to potential insecticide selection pressure on the vector. With the regular use of insecticides, it is important to monitor the development of resistance patterns at the earliest possible stage to assess their likely impact on control operations [13]. Comprehensive and regular susceptibility testing is therefore required to form an essential adjunct to any insecticide-based control operation. To ensure the insecticide-based interventions are cost-effective, an insecticide must be selected to which the mosquito vectors are susceptible.

This study investigated the insecticide susceptibility or resistance status in dengue vectors from the National Capital Region (NCR) based on the standard World Health Organization (WHO) contact test using diagnostic doses. Determining the resistance in the mosquito vectors will provide essential information for health planners and disease control specialists in selecting alternative compounds and in modifying vector control strategies currently implemented. Resistance data should also propel more studies on the mechanisms of resistance.

## Materials and Methods

### Mosquito collections

The field-caught strains of *Ae. aegypti* used in this study were collected in the National Capital Region (NCR), also known as Metropolitan Manila, located in the southwestern portion of Luzon (Figure 1). NCR is the smallest administrative region (620 km^2^) but is the second most populous region with approximately 13.5 million inhabitants [14]. It is comprised of 16 cities and one municipality grouped into four districts: Capital District, Eastern Manila, Northern Manila and Southern Manila. In each city/municipality, three barangays (brgy.) were selected as sites for mosquito collection (S1 Table) based on the current and previously recorded dengue cases and the City Health Offices’ recommendations.

**Figure 1.**
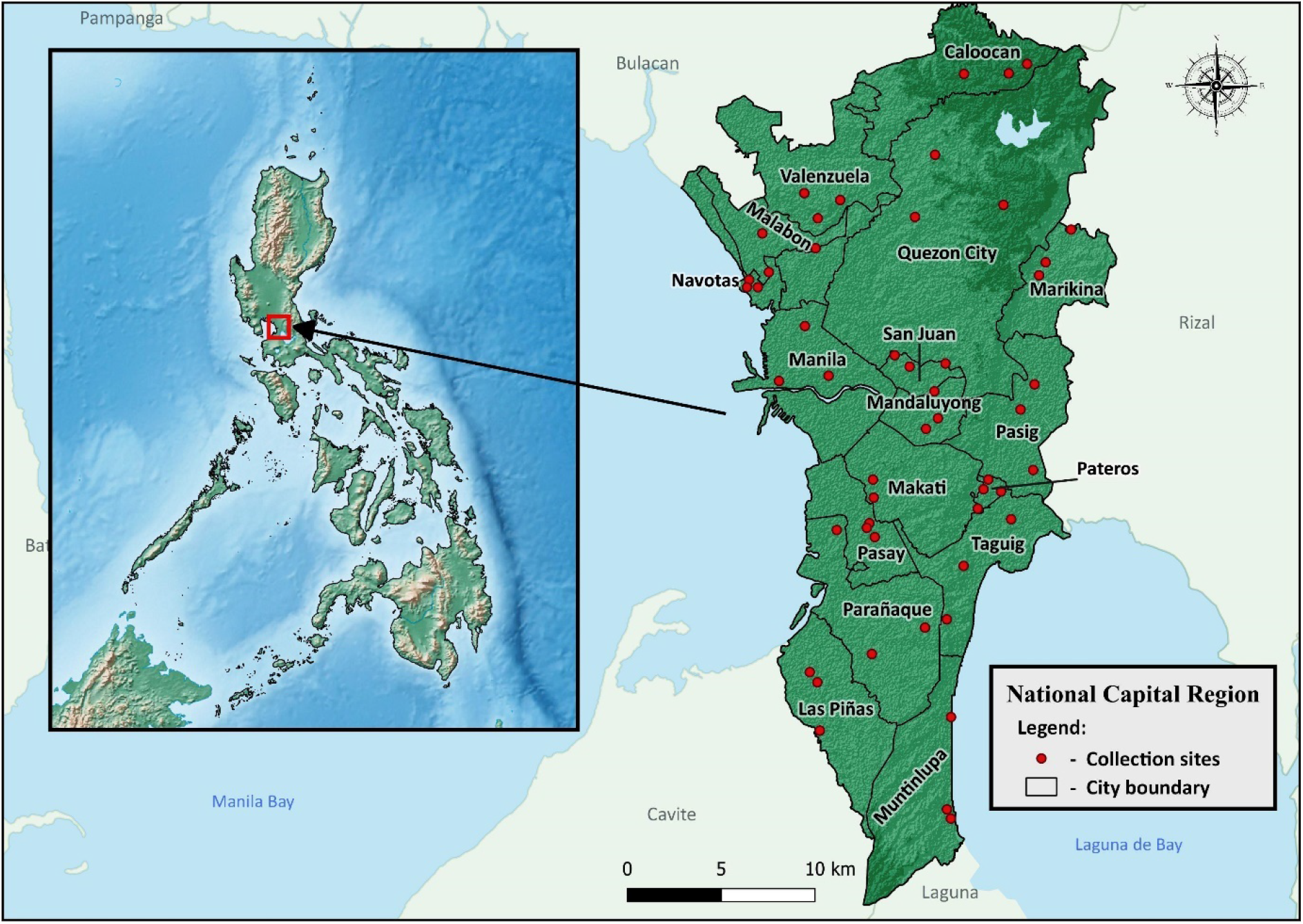
Location of *Aedes* mosquito collections in each city/municipality of the National Capital Region

Eggs of *Aedes* mosquitoes were collected between September 2013 and June 2015 using oviposition traps (ovitraps). The trap consists of 300ml plastic pots, two-thirds filled with water, and a wooden stick (paddles) made of high-density fiberboard wrapped with filter paper was added as an oviposition medium. The ovitraps were installed in 30 houses per barangay; placed indoors and outdoors. The wooden paddles were collected after one week and placed in labeled plastic bags. Emerged larvae in the traps were also collected and placed in separate containers. Immature stages (larva and pupa) of mosquitoes were also collected from other water-holding containers within the vicinity to augment ovitrap collections.

All collected samples were brought to the laboratory and reared in an insectary at 26±2°C and 60±10% relative humidity. Emerged adult mosquitoes were identified morphologically and species from same barangay was pooled in the same cage. The larvae were fed with ground fish flakes (Tetrafin®) and Brewer’s yeast while adults were provided with 10% sugar solution *ad libitum*. Female mosquitoes were given blood meals using white mice to induce egg laying. Rearing of the mosquitoes was continued until the F_1_ and F_2_ generations.

### Insecticide Susceptibility Tests

Tube bioassays were carried out using standard insecticide-treated papers following the WHO guidelines [15-17]. Insecticide-impregnated and oil-impregnated control papers were sourced from Vector Control Research Unit, Universiti Sains Malaysia; a WHO Collaborating Center. The following diagnostic concentrations of insecticides were used: pyrethroid (0.75% permethrin, 0.05% deltamethrin, 0.15% cyfluthrin, 0.05% lambdacyhalothrin, 0.05% etofenprox), organochlorine (4% DDT) and organophosphate (5% malathion). Diagnostic doses recommended for *Aedes* in the WHO adult bioassays have not yet been defined for some insecticides; thus, *Anopheles* mosquito’s listed diagnostic doses were used in this study. These doses are higher than *Aedes’* for the few insecticides defined in 1992 [17] and were also used in similar published studies [18-21].

For each insecticide, five batches of 20 non-bloodfed females (5-7 day old) were introduced into holding tubes lined with untreated paper for 60 minutes for the exposed group while three batches of 20 mosquitoes were provided as the control group. Mosquitoes were then transferred into exposure tubes lined with insecticide-treated or control paper for 1 hour. In this study, modification of the WHO procedure (i.e. recording of knock down every 5 minutes interval) was made to develop a sensitive detection method for monitoring resistance trends [22,23]. After exposure, mosquitoes were transferred back to the holding tubes and had *ad libitum* access to sugar solution. Mortality was determined 24 hours post-exposure in each replicate.

### Data Analysis

The mortality rate for each insecticide was calculated by adding the number of dead mosquitoes on all five exposure replicates and expressing as a percentage of the total number of exposed mosquitoes. A similar calculation was made for the control mortality. When control mortality was between 5 and 20%, mortality was corrected using Abbott’s formula [24]; with negative values rounded to zero.

Following the WHO guidelines [16], mosquito populations were considered susceptible if the percentage of mortality ranges from 98 to 100% at 24 hours post-exposure. Mosquito mortality between 90 and 97% was defined as incipient or developing resistance but would require follow-up testing to confirm resistance. If mortality is less than 90%, phenotypic resistance is confirmed if at least 100 mosquitoes were tested.

## Results

Ovitraps in the 51 barangays were able to collect immatures of *Ae. aegypti, Ae. albopictus* and *Culex quinquefasciatus*; the majority of which were *Ae. aegypti*. Due to the insufficient test specimen for *Ae. albopictus*, bioassays were done on *Ae. aegypti* from 31 barangays only. Mortality and knockdown (KD) rates for each mosquito strain are reported in S2 Table and only those tests with at least 100 mosquitoes in all replicates were included in the final analysis.

Mosquito populations from the different barangays showed varying mortality rates based on the WHO susceptibility tests conducted against different insecticides. Resistance to permethrin and etofenprox was detected in all tested *Ae. aegypti* populations with mortality rates ranging from 1.00% to 86.91% (mean = 52.28 ± 26.49) and 0.00% to 40.00% (mean = 14.34 ± 11.38) for permethrin and etofenprox, respectively. Exposure to deltamethrin yielded mortality rates ranging from 41.00% to 100.00% (mean = 85.11 ± 20.38) indicating varying resistance levels in the *Ae. aegypti* populations. More than half of the populations (10 out of 18) showed incipient resistance to deltamethrin while five populations had confirmed phenotypic resistance. Only *Ae. aegypti* populations from barangays 183 (Manila), Putatan (Muntinlupa City) and Tandang Sora (Quezon City) showed susceptibility to deltamethrin. Varying resistance levels to cyfluthrin were also detected in tested populations with mortality rates from 50.00% to 100.00% (mean = 82.17 ± 17.58); with five populations with confirmed resistance, another five populations showing incipient resistance while only *Ae. aegypti* population from Brgy. Cupang (Muntinlupa City) showing full susceptibility. Exposure to lambdacyhalothrin rendered mortality rates ranging from 16.00% to 93.00% (mean = 69.35 ± 22.85) showing confirmed resistance in nine populations while three populations with incipient resistance. Phenotypic resistance to DDT was observed in all tested populations with test mortality ranging from 2.00% to 47.00% (mean = 15.44 ± 12.04). In contrast, all *Ae. aegypti* populations tested (n=13) against malathion were susceptible to the insecticide with 100% test mortality in all tests.

### Knockdown rates

Overall knockdown and mortality rates for the NCR mosquito population are reported in Figure 2 while evolution of the KD rate over a 60-minute observation period is shown in Figure 3. Overall KD rate for cyfluthrin (90.73%) and deltamethrin (86.28%) is higher compared to mortality (82.17% and 85.11%, respectively) showing recovery in the mosquito population during the 24-hour recovery period. Higher mortality rates were observed for permethrin, lambdacyhalothrin, etofenprox, DDT and malathion compared to KD rate.

**Figure 2.**
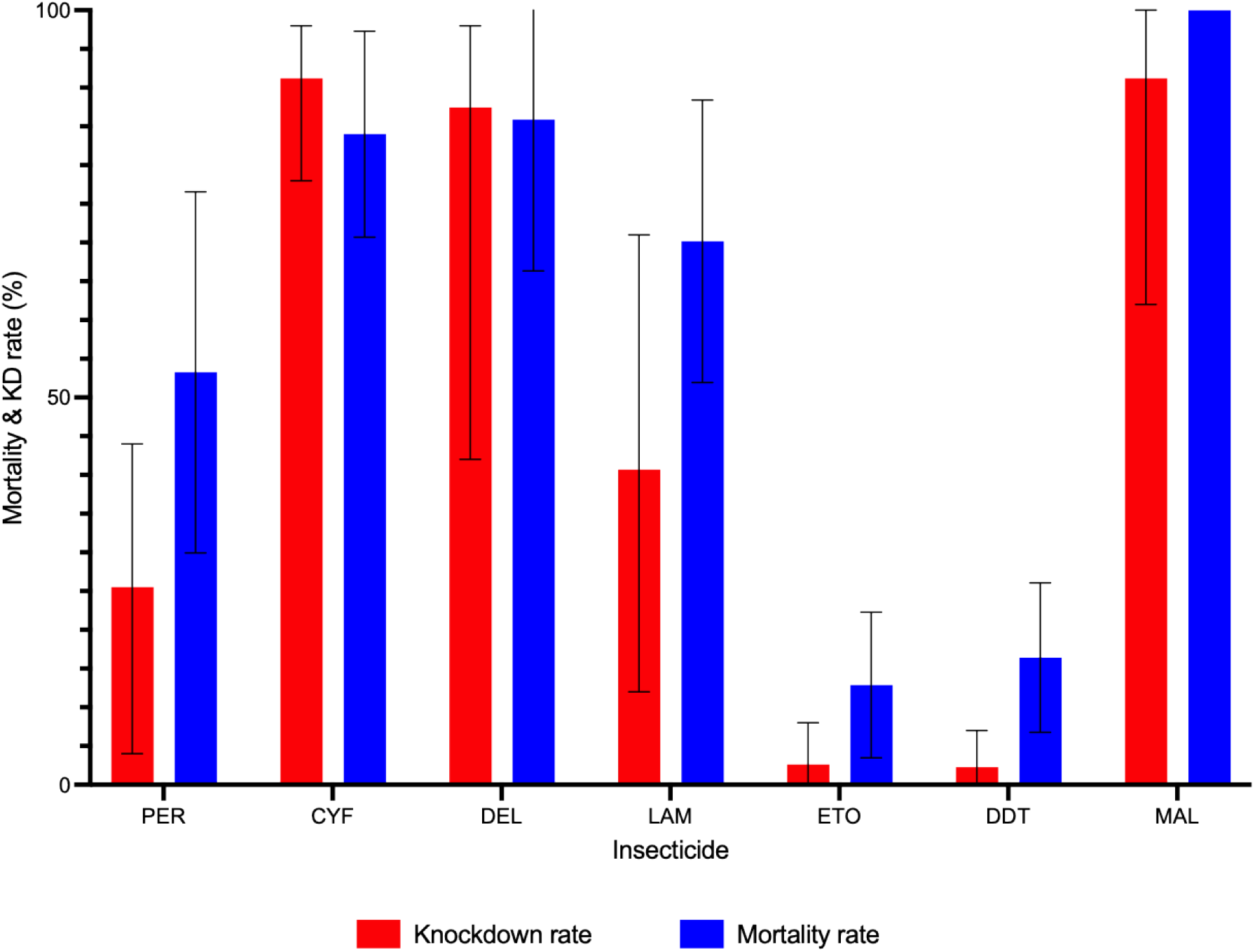
Mortality and knockdown rate determined following the WHO susceptibility test procedure. CYF – cyfluthrin, DEL – deltamethrin, ETO – etofenprox, LAM – lambdacyhalothrin, MAL – malathion, PER – permethrin, error bar – standard deviation

**Figure 3.**
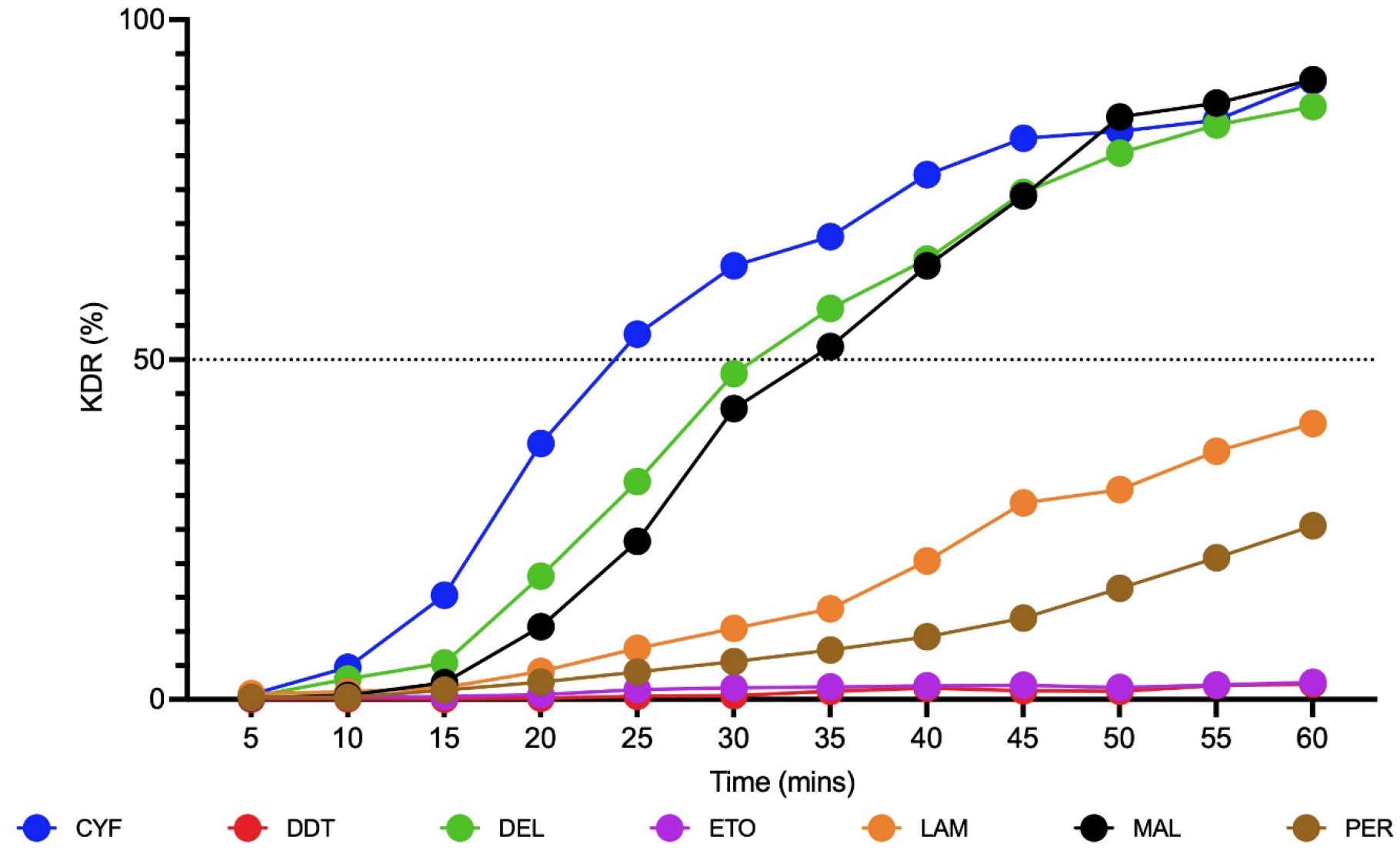
Evolution of the knockdown rate during insecticide exposure recorded every five minutes during the exposure period to insecticide. Dash line indicates a 50% KD rate

## Discussion

Here, we provide evidence of the resistance status of *Ae. aegypti* populations from NCR, Philippines to insecticides used to control adult mosquitoes. *Aedes aegypti* populations from this region exhibited varying susceptibility/resistance levels to pyrethroid insecticides, confirmed resistance to DDT and susceptibility to malathion. To the authors’ knowledge, this is the first published report to establish the insecticide resistance of *Ae. aegypti* populations from the Philippines.

There had been limited studies on insecticide resistance in mosquito vectors from the Philippines; most of these were focused on the malaria vectors [22,23,25-28]. Early studies reported that *Ae. aegypti* from Clark Air Base in Pampanga is susceptible to DDT and dieldrin [27]. In contrast, the secondary dengue vector *Ae. albopictus* showed resistance to BHC/cyclodienes [26]. From 2005 to 2006, a thesis study detected resistance in *Ae. Aegypti* populations from Manila to permethrin, cyfluthrin, deltamethrin and DDT but were still susceptible to malathion and temephos [29]. In the same study, populations from Cabuyao, Laguna exhibited varying resistance/susceptibility to the said insecticides.

Available guidelines on adulticide application of malathion, fenitrothion or permethrin using ULV mist blowers during dengue epidemics can be traced back to 1991 [30,31]. From 1997 to 2003, the local government units invariably used permethrin, deltamethrin, cyfluthrin and organophosphates such as pirimiphos-methyl and malathion. From 2004 onwards, other insecticides registered with the Fertilizer and Pesticide Authority (FPA) were used for space spraying while permethrin-treated curtains were included as an additional adult control measure [32]. The use of insecticides inevitably induces selection pressures on the target vectors; particularly in the primary vector of dengue *Ae. aegypti*.

Pyrethroids were heavily used to control dengue vectors due to their immediate result, high mortality effect, low residual efficacy and low mammalian toxicity [33-35]. Based on DOH regional offices’ reports, activities involving chemical control like thermal fogging, space spraying, and insecticide-treated curtains were all leaning towards the use of pyrethroids. With the high dependence, frequency and usage of pyrethroid-based insecticides, higher selection pressure to the insecticides could have resulted in increased resistance levels in mosquitoes [36,37].

High resistance to etofenprox was detected in all tested populations using the recommended diagnostic dose for *Aedes*; with the highest test mortality of only 40%. On the other hand, some level of resistance to cyfluthrin was detected in *Ae. aegypti* populations except those from Brgy. Cupang (Muntinlupa). Similarly, cyfluthrin resistance was also observed in *Ae. aegypti* from Jakarta, Indonesia [38].

The *Anopheles* diagnostic doses were used for permethrin, deltamethrin and lambdacyhalothrin since no defined diagnostic doses for *Aedes* were recommended when this study was conducted. Despite the higher *Anopheles* diagnostic doses used, resistance was observed to Type I (permethrin) and Type II (deltamethrin, lambdacyhalothrin) pyrethroids. Similar findings were observed in *Ae. aegypti* populations from neighboring Southeast Asian countries, particularly the observed resistance to permethrin [7,38-43]. All tested populations also showed some level of resistance to lambdacyhalothrin; most of the tested populations with confirmed phenotypic resistance while the remaining populations showed incipient resistance; similar to observations in Indonesia [38]. Varying levels of resistance to deltamethrin were detected in the different *Ae. aegypti* populations; with majority showed tolerance to deltamethrin. Compared with *Ae. aegypti* from Malaysia, all tested populations were already resistant to deltamethrin with test mortality ranging from 0 to 82% [43]. In contrast, *Ae. aegypti* populations from Laos are mostly susceptible to the insecticide [44].

DDT has long been banned in the country [30,45] yet resistance was observed in the mosquito populations to the insecticide. Pyrethroids and organochlorines have the same mode of action that interrupts nerve impulses in nerve axons that causes hyperexcitation and tremor followed by paralysis and blocking of nerves in mosquitoes [7,46]. Resistance of the *Ae. aegypti* populations to pyrethroids may have conferred the populations’ resistance to DDT. However, further studies are needed to establish the possible cross-resistance to the two insecticide classes. Mutations in resistance against permethrin and deltamethrin had already been reported to be linked to DDT resistance, showing that there can be cases of cross-resistance among the two insecticide groups [47,48]. DDT’s persistence in the environment is also considered as one of the causes of resistance [49,50].

Organophosphates have been heavily used for dengue vector control and in agriculture, leading to intense selection pressure and highly resistant mosquitoes in other countries [51]. In the Philippines, organophosphates such as malathion are often used against agricultural pests but was also used against dengue vectors by the different local government units [30]. The regular use of malathion in an urban setting is a challenge due to the insecticide’s repulsive odor; thus, it may explain *Ae. aegypti* populations’ susceptibility to the insecticide. However, regular resistance monitoring for this insecticide is needed as resistance to malathion has already been reported in other neighboring countries [7,38,52]. A more detailed retrospective study to determine the extent and manner from which the compounds were used for vector control purposes; either as part of the disease control program or as part of personal protection measures; should be done including insecticide-related practices that would impact the judicious use and environmental safety. This study should therefore impact the current DOH policy and the advocacy of the National *Aedes*-borne Viral Disease Prevention and Control Program.

Results for some insecticides in this study used the higher diagnostic doses recommended for *Anopheles* compared to the recently published recommended dose for *Ae. aegypti* [15-17,53]. Thus, the data presented herein may not show the field-caught *Aedes aegypti* populations’ current level of susceptibility. Also, the results presented pertain to standardized laboratory bioassays following existing guidelines and using suggested diagnostic dosages. Susceptibility in such conditions does not necessarily reflect susceptibility or resistance level translated into the efficacy of the programmatically-applied interventions in the field. Further studies of insecticide efficacy should be conducted in areas with known and characterized resistance levels to confirm field efficacy.

Results from this study focused on phenotypic resistance in *Ae. aegypti* populations in NCR, Philippines. Nonetheless, this study provided pertinent information on *Ae. aegypti’s* general susceptibility or resistance to insecticides and serves as baseline data for future studies on insecticide resistance in dengue vectors and guides public health authorities and program managers. Moreover, this report should signal an update on the vector control program’s current insecticide resistance management plan. Further studies on the characterization of resistance mechanisms will help provide the impetus for developing newer compounds, continued studies on mosquito resistance’s molecular/genetic basis and the possibility of genetic manipulation to maintain susceptibility in vector populations.

There is a need to assess the operational impact of pyrethroid use in controlling adult mosquitoes. The use of compounds with a different mode of action from pyrethroids is highly recommended. Since the current formulation of malathion renders it unacceptable to the community, reformulation of the product by the manufacturers may help increase acceptance and use in dengue vector control. In the meantime, the Dengue Control Program’s 4S advocacy is highly encouraged for disease prevention and control: (1) Search and destroy mosquito breeding sites, (2) Secure self-protection, (3) Seek early consultation, and (4) Support fogging/spraying only in hotspot areas where increase in cases is registered for two consecutive weeks to prevent an impending outbreak.

## Supporting information

S1 Table. GPS coordinates of mosquito collection sites in the National Capital Region, Philippines

S2 Table. Knockdown and mortality rates of adult *Aedes aegypti* in the susceptibility test using the WHO standard bioassay

## Acknowledgements

The authors would like to thank the RITM Medical Entomology staff for assisting with the mosquito collection, rearing and bioassays and Mr. Jonathan Sabellano for creating the map. We thank DOH-NCR Office and the city/municipal health offices for their support and cooperation in the study.

## Author Contributions

**Conceptualization:** Ferdinand Salazar, Jason Angeles

**Methodology:** Ferdinand Salazar, Jason Angeles, Richard Paul Malijan

**Investigation:** Jason Angeles, Ariza Minelle Apilado, Mary Ann Ammugauan

**Supervision:** Ferdinand Salazar

**Data Curation:** Richard Paul Malijan, Jason Angeles

**Formal Analysis:** Richard Paul Malijan

**Writing – Original Draft Preparation:** Jason Angeles, Richard Paul Malijan

**Writing – Review & Editing:** Richard Paul Malijan, Ferdinand Salazar, Ariza Minelle Apilado

